# Small Molecule Degradation of the microRNA-21 Precursor Rescues Pathogenic Pathways in Cellular Models of Fibrosis

**DOI:** 10.64898/2025.12.09.693244

**Authors:** Tenghui Wang, Xueyi Yang, Yeongju Lee, Jin H. Song, Jessica L. Childs-Disney, Joe G.N. Garcia, Matthew D. Disney

## Abstract

MicroRNAs (miRNAs) are short RNA molecules that bind to target mRNAs, resulting in translational repression and gene silencing. Overexpression of microRNA-21 (miR-21) is associated with various human diseases, including autosomal dominant polycystic kidney disease (ADPKD) and pulmonary fibrosis. In this study, a previously described heterobifunctional molecule, TGP-21-RiboTAC, that degrades the miR-21 precursor (pre-miR-21) in triple negative breast cancer cells was investigated in polycystic kidney cell lines and a lung fibroblast cell line. In the former, TGP-21-RiboTAC degraded pre-miR-21 and de-repressed of miR-21’s downstream target, Programmed Cell Death 4 (PDCD4) and Peroxisome Proliferator-Activated Receptor alpha (PPARα), known drivers of ADPKD. The heterobifunctional molecule also inhibited cyst growth and rescued the metabolic alterations that occur in ADPKD. In the lung fibroblast cell line, MRC-5, TGP-21-RiboTAC also reduced pre- and mature miR-21 levels, rescued Transforming Growth Factor β (TGF-β)-induced repression of SMAD Family Member 7 (SMAD7) and inhibited cell invasion. Collectively, these studies demonstrate the potential of targeted RNA degradation as therapeutic agents that retard the development of organ fibrosis.

## INTRODUCTION

Micro (mi)RNAs are small, non-coding RNAs that are 20-23 nucleotides long and control gene expression by binding to the 3’ untranslated region (UTR) of complementary mRNAs, causing degradation or translational repression^1^. As miRNAs control essential cellular processes, mutation and/or aberrant miRNA expression is associated with a variety of pathobiologies. Thus, removal of overexpressed disease-causing miRNAs is a viable therapeutic strategy alleviate disease severity.

Various studies have linked microRNA-21 (miR-21) with pulmonary and kidney fibrosis^2-4^. Like other miRNAs, miR-21’s biogenesis involves two sequential processing steps: conversion of the primary (pri-) miRNA to its precursor (pre-) form in the nucleus by Drosha and conversion of the pre-miRNA to the mature, active miRNA in the cytoplasm by Dicer^5^ (**Fig. 1A**).

**Figure 1:**
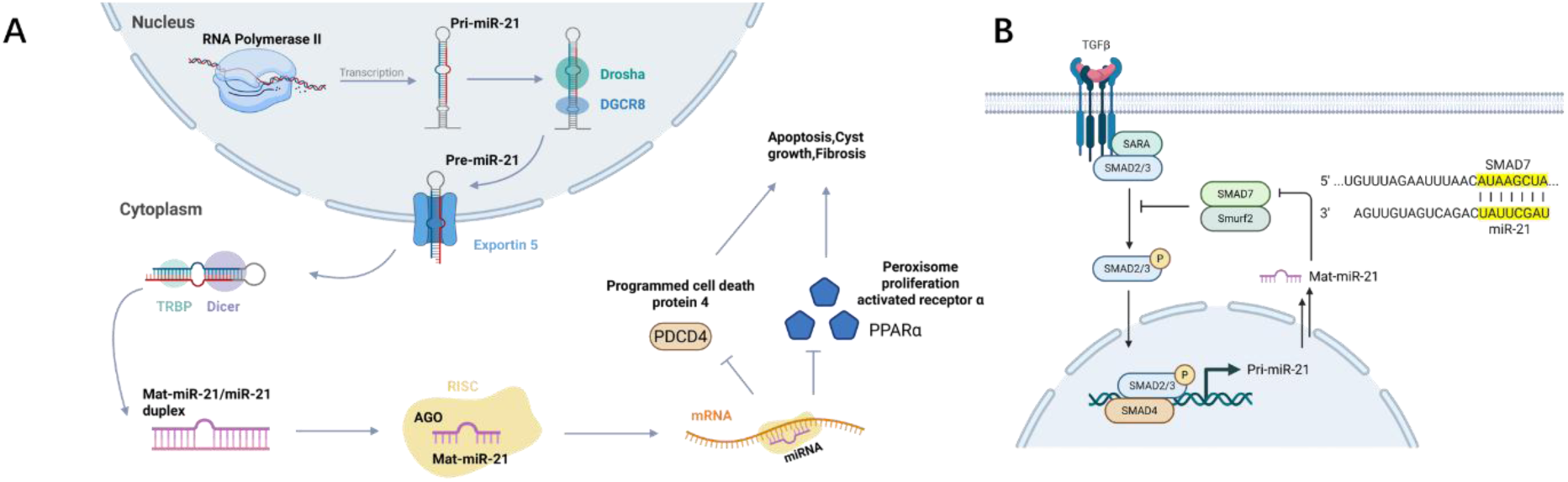
miR-21-regulated pathway in autosomal dominant polycystic kidney disease (ADPKD) and pulmonary fibrosis. **A,** Schematic representation of miR-21–mediated inhibition of PDCD4 and PPARα that promotes cyst growth and fibrosis.**B,** Schematic represnetation of miR-21 represses SMAD7, amplifying the Transforming Growth Factor Beta (TGF-β) signaling pathway.

Autosomal dominant polycystic kidney disease (ADPKD) is a hereditary disorder characterized by the formation of fluid-filled renal cysts, associated with kidney enlargement and progressive renal fibrosis^6^, often resulting in renal failure, cardiovascular complications, and associated morbidities^6-8^. The role of miR-21 ^8^ in ADPKD and the development of fibrosis has been validated in vivo where inactivation of miR-21 by an antagomir oligonucleotide prevents fibrosis while overexpression enhances renal fibrosis ^9-11^. MiR-21 mediates its pathogenic effects by repressing expression of Programmed Cell Death 4 (PDCD4), a protein which suppresses apoptosis in cyst epithelial cells by inhibiting the translation of procaspase-3 mRNA^12, 13^ to promote cyst growth^4^. MiR-21 also reduces expression of Peroxisome Proliferator-Activated Receptor alpha (PPARα)^4, 14^ that regulates genes involved in various aspects of lipid metabolism including fatty acid uptake, utilization, synthesis and degradation^15^. miR-21-mediated PPARα repression reduces fatty ^7^acid oxidation (FAO) which promotes cyst growth^16^.

Aberrant miR-21 expression has also been implicated in idiopathic pulmonary fibrosis^9, 17, 18^, the most common form of interstitial lung disease^19^ presumably via amplification of Transforming Growth Factor Beta (TGF-β) signaling pathway known to be intimately involved in development of fibrotic lung diseases. Upregulation of miR-21 by TGF-β suppresses expression of SMAD Family Member 7 (SMAD7), thereby driving fibrosis progression **(Fig. 1B)**. As the miR-21 KO mouse is without an adverse phenotype^4^, inhibiting miR-21 biogenesis is a potential therapeutic strategy to ameliorate both kidney and lung fibrosis where miR-21 serves as a disease driver.

One approach to mediate RNA decay is to exploit ribonucleases (RNases) that naturally regulate RNA lifespan and recruit them to specific transcripts via a heterobifunctional small molecule, or ribonuclease targeting chimera (**RiboTAC**)^20^ representing the small molecule equivalent of antisense oligonucleotides. We previously reported that **TGP-21-RiboTAC** selectively induces cleavage of oncogenic pre-miR-21 in triple-negative breast cancer cells and in a xenograft mouse model by binding to the RNA target and inducing proximity of RNase L to effect targeted degradation^21^. RNase L is an integral part of the viral immune response and is present in minute quantities in all cells as an inactive monomer^22^. To address the suitability of **TGP-21-RiboTAC** targeting to mitigate miR-21-driven diseases, we evaluated for targeted degradation of pre-miR-21 in in vitro cellular models of ADPKD and pulmonary fibrosis. This study illustrates the potential applicability of **TGP-21-RiboTAC** reduce both pre-and mature miR-21 levels, de-repressed downstream proteins, and improved ADPKD-associated and lung fibrotic cellular phenotypes. Thus, the study illustrates the potential applicability of **TGP-21-RiboTAC** represents a novel therapeutic approach to diverse fibrotic disorders with limited treatment options.

## RESULTS

### TGP-21-RiboTAC reduces pre-miR-21 and mature miR-21 levels in ADPKD cells in RNase L-dependent manner

**TGP-21-RiboTAC** comprises a dimer that binds an A and a U bulge in and around the Dicer site of pre-miR-21 simultaneously (named **TGP-21**; *K*d = 1 ± 0.1 µM^23^) and an RNase L recruiter dubbed **C1-3** (**Fig. 2B**). To confirm whether the heterobifunctional molecule degrades pre-miR-21 in the context of kidney fibrosis, two cell lines were employed, the mouse polycystic kidney cell line, mIMCD-3, and the human polycystic kidney cell line, WT-9-7. Importantly, the **TGP-21** binding site, including the Dicer processing site and apical loop, is conserved in human and mouse **(Supplementary Fig. 1A)**.

**Figure 2:**
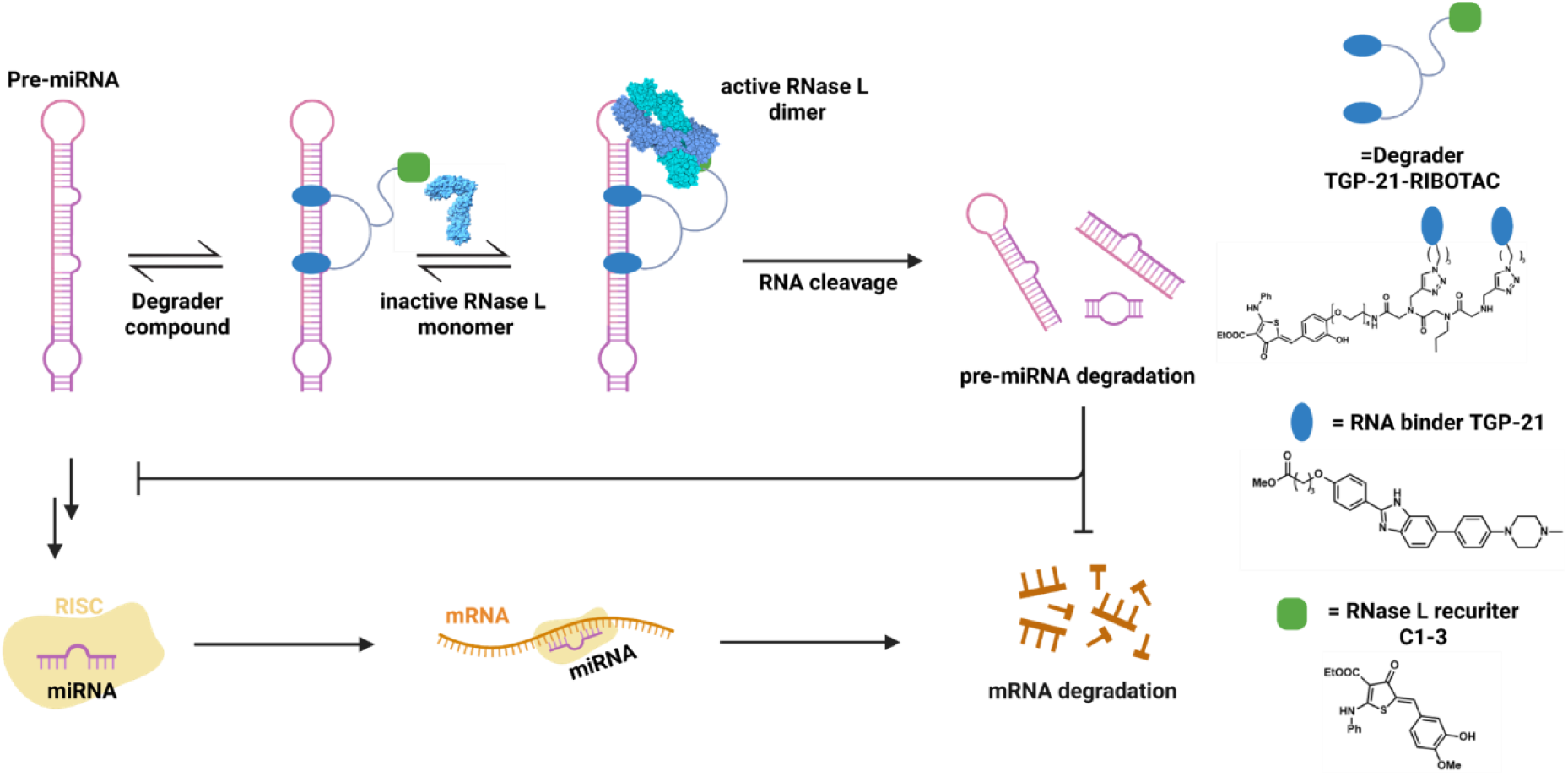
Mechanism of degradation of pre-miRNAs induced by RiboTACs. RiboTACs bind to structures in miRNA precursors (pri- or pre-) and recruits endogenous RNase L to cleave the target by induced proximity. Degradation of pre-miRNA prevents expression of the mature miRNA and thus de-represses downstream mRNA targets. Structure of degrader **TGP-21-RiboTAC**, RNA binder **TGP-21,** and RNase L recruiter **C1-3** are shown on the right side.

To determine the concentration range suitable for these studies, the viability of both cell lines was measured as a function of **TGP-21-RiboTAC** concentration (**Supplementary Fig. 2A-B**). These results showed that concentrations less than 5 µM are well tolerated, and thus 1 µM was chosen as the highest concentration to avoid spurious results that might be associated with toxicity. Dose dependent reduction of pre-miR-21 by **TGP-21-RiboTAC** was observed in mIMCD-3 cells, with statistically significant reduction of 45±8% (p < 0.05) observed at the 1 μM dose (**Fig. 3A**). Likewise, miR-21-5p abundance was reduced dose dependently with a maximal reduction of 62±17% (p < 0.001; **Fig. 3B**). **TGP-21-RiboTAC** was less potent in WT-9-7 cells where reduction was only observed at the 1 μM dose pre-miR-21 by 28±7% (p < 0.05); dose dependent reduction of miR-21-5p was observed, however (55±17% (p < 0.001) upon treatment with 1 μM of **TGP-21-RiboTAC** (**Fig. 3A-B**).

**Figure 3:**
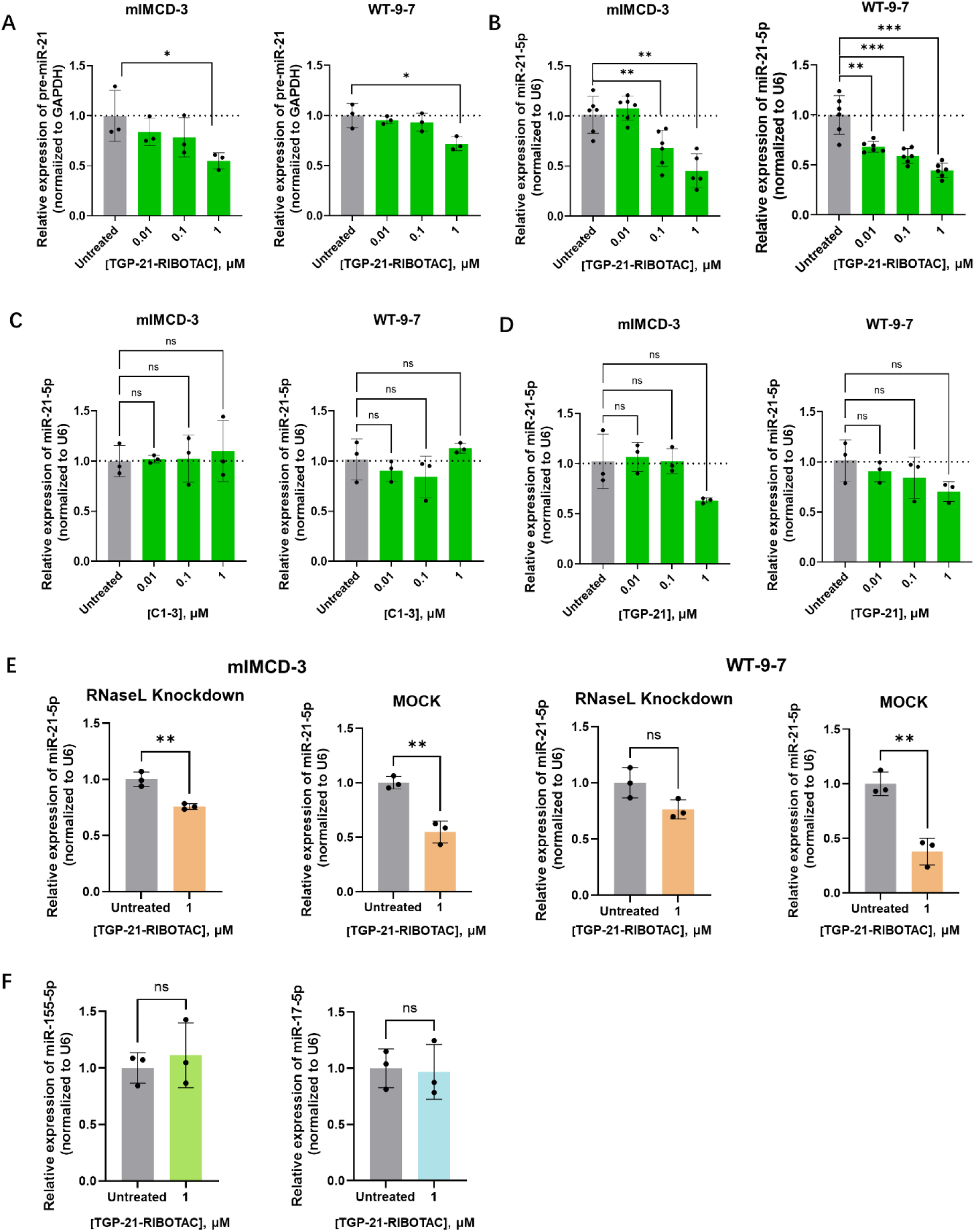
Evaluation of TGP-21-RiboTAC bioactivity in polycystic kidney cell lines. **A,** Effect of **TGP-21-RiboTAC** on pre-miR-21 levels in mIMCD-3 and WT-9-7 cell lines, as determined by RT-qPCR (n = 3 biological replicates). **B,** Effect of **TGP-21-RiboTAC** on miR-21-5p abundance in mIMCD-3 and WT-9-7 cell lines, as determined by RT-qPCR (n = 6 biological replicates per treatment group). **C,** Effect of **TGP-21-RiboTAC** on miR-21-5p levels in WT-9-7 cell line where RNase L was knocked down with an siRNA, as determined by RT-qPCR (n = 2 biological replicates for untreated; n = 3 biological replicates for **TGP-21-RiboTAC**-treated samples). **D,** Effect of **TGP-21-RiboTAC** on miR-155-5p and miR-17-5p levels in WT-9-7 cells, as determined by RT-qPCR (n = 3 biological replicates). **E,** Effect of RNase L recruiter **C1-3** on miR-21-5p levels in mIMCD-3 and WT-9-7 cells. **F,** Effect of pre-miR-21 binder **TGP-21** on miR-215p abundance in mIMCD-3 and WT-9-7 cells. p < 0.05; **, p < 0.01; ***, p < 0.001; and ****, p < 0.0001, as determined by a One-way ANOVA with multiple comparisons (panels A and B) or t test (panels C and D). Data are reported as mean ± SD.

The effects of pre-miR-21 binder **TGP-21** and the RNase L recruiter **C1-3** on miR-21-5p levels were measured in mIMCD-3 and WT-9-7 cell line **(Fig. 3C-D**). **TGP-21** was previously shown to reduce miR-21-5p, but not pre-miR-21 abundance in triple negative breast cancer cells^21^. No statistically significant effect was observed for RNase L recruiter **C1-3** in either cell line at any concentration tested (0.01, 0.1 and 1 µM; **Fig. 3C**). As expected, reduction of miR-21-5p levels were observed upon treatment of mIMCD-3 and WT-9-7 cell with 1 µM of **TGP-21** (**Fig. 3D**).

It was further validated that the reduction of pre- and mature miR-21-5p abundance is RNase L-dependent by treating cells with an RNase L-targeting siRNA pool. The activity of **TGP-21-RiboTAC** for reducing miR-21-5p levels was attenuated upon RNase L knock down. In mIMCD-3 cells, activity was reduced from 46±7% (p<0.01) reduction to 24±4% (p<0.01) reduction, while in WT-9-7, miR-21-5p levels were only reduced to 24±10% (no significance) upon RNase L knock down (as compared to 62±9% (p<0.01)) **(Fig. 3E)**. Collectively, the activity data for **TGP-21-RiboTAC, TGP-21**, and **C1-3** as well as the RNase L-dependency studies support that the RiboTAC binds to pre-miR-21 and recruits RNase L to induce the RNA’s degradation.

The selectivity of **TGP-21-RiboTAC** was previously extensively studied in MDA-MB-231 cell line across the miR-nome, amongst highly abundant transcripts, and the proteome^21^.To provide some insight into the selectivity of **TGP-21-RiboTAC** in WT-9-7 cells, its effect on two other miRNA associated with ADPKD and other kidney diseases, miR-17 and miR-155^24-26^, was measured. As the secondary structures of these two miRNAs are very different from miR-21 (**Supplementary Fig. 1B**), it is hypothesized that **TGP-21-RiboTAC** would not affect their abundance. Indeed, RT-qPCR analysis revealed that neither miR-17-5p nor miR-155-5p abundance was affected upon treatment of WT-9-7 cells with 1 µM of **TGP-21-RiboTAC** (**Fig. 3F**). These data suggest that the **TGP-21-RiboTAC** is specific for miR-21 and that de-repression of its downstream targets and hence deactivation of the miR-21-ADPKD circuit, *vide infra*, is an on-target effect.

### Reduction of miR-21 by TGP-21-RiboTAC derepresses PPARα and PDCD4

As aforementioned, PPARα, a direct target of miR-21^11, 14^, is a transcription factor that regulates many aspects of cellular metabolism^15, 25^ (**Fig. 4A**). Genetic studies demonstrate that deletion of *pparα* aggravates cyst growth in a slowly progressive mouse model of ADPKD^16, 25^. Therefore, the effect of **TGP-21-RiboTAC** on both PPARα protein levels, its downstream targets, and metabolism were evaluated.

**Figure 4:**
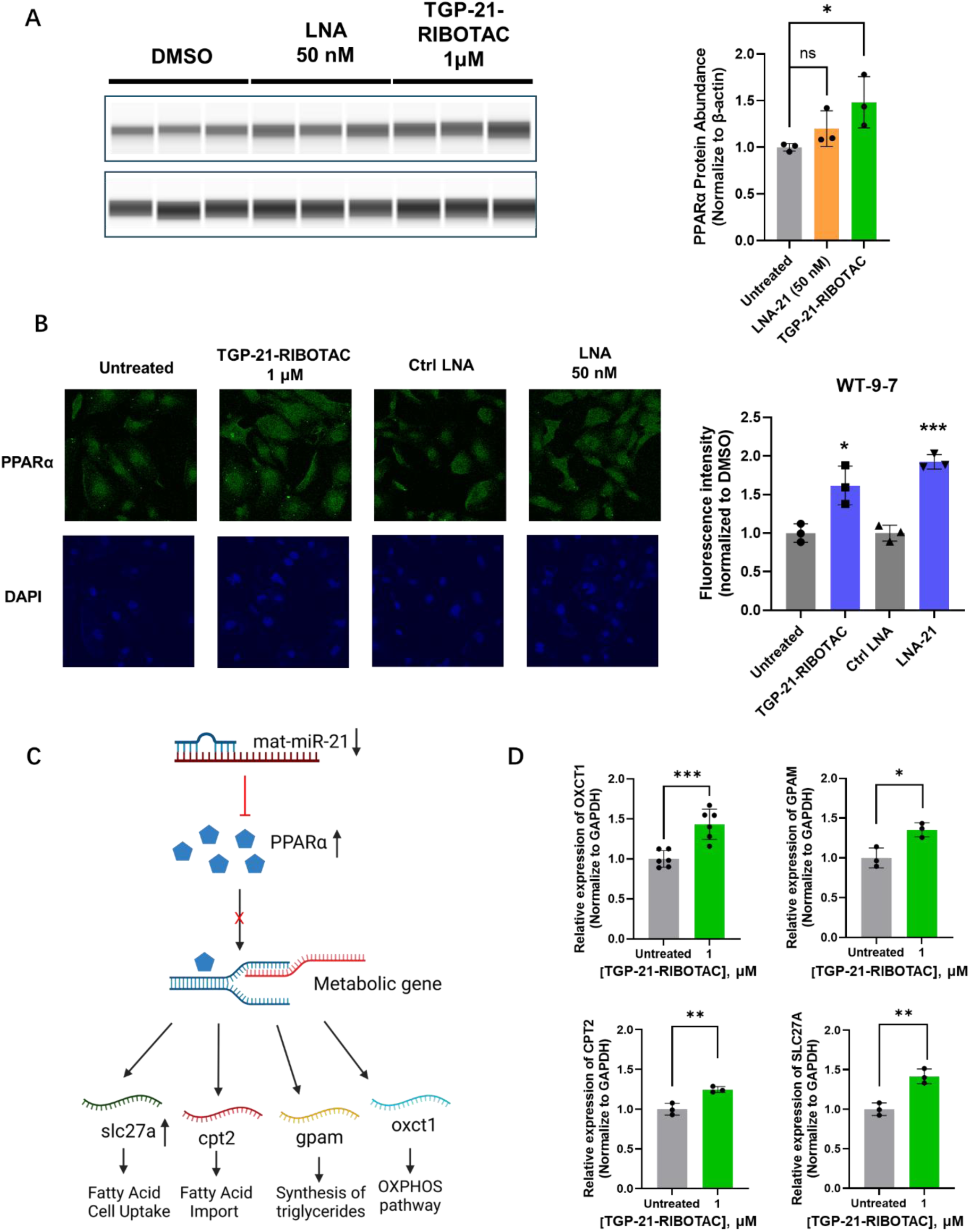
TGP-21-RiboTAC de-represses PPARα and the transcription factor’s downstream targets in polycystic kidney cells. **A,** PPARα protein expression in mIMCD-3 cell line upon treatment with **TGP-21-RiboTAC**, as assessed by Simple western (n = 3 biological replicates). **B,** Representive images and quantification of PPARα proein expression in WT-9-7 cell line upon **TGP-21-RiboTAC** treatment, as assessed by immunofluorescence (IF) imaging (n = 3 biological replicates). **C,** Schematic representation of miR-21–mediated repression of **PPARα** amd its downstream target genes (*slc27a*, *cpt2*, *gpam*, *oxct1*). **D,** Effect of **TGP-21-RiboTAC** on PPARα target genes *oxct1*, *gpam*, *cpt2* and *slc27a* expression in WT-9-7 cells, as determined by RT-qPCR (n = 3 biological replicates). *, p < 0.05; **, p < 0.01; ***, p < 0.001; and ****, p < 0.0001, as determined by t tests. Data are reported as mean ± SD.

The effect of 1 μM **TGP-21-RiboTAC** treatment on PPARα protein levels in mIMCD-3 cells and WT-9-7 cells were assessed by both immunofluorescence (IF) and Western blotting. A 48±27% (p < 0.05) increase of the protein was observed in mIMCD-3 cells and 61±25% (P < 0.001) increase in WT-9-7 cells (**Fig. 4A-B**). These results illustrate that a decrease of miR-21-5p levels induced by 1 μM of ∼50% is sufficient to de-repress PPARα expression. No effect was observed at the *PPARα* transcript level in either cell lines (**Supplementary Fig. 3A-B**).

To determine if the observed increases in PPARα protein was sufficient to upregulate its downstream targets, transcript levels for 3-oxoacid CoA-transferase 1 (OXCT1)^25^, fatty acid transport proteins (SLC27A)^25, 27^, carnitine palmitoyl-transferase 2 (CPT2)^25, 27^, and glycerol-3-phosphate acyltransferase (GPAM)^25^ were measured in WT-9-7 cells by RT-qPCR. At a dose of 1 µM of **TGP-21-RiboTAC**, the abundance of each transcripts was increased, by 43±19% (p < 0.001), 41±9% (p < 0.05), 24±4% (p < 0.05) and 35±9% (p < 0.05), respectively (**Fig. 4D**), similar effect was observed for OXCT1 ( 43±9%, (p < 0.001)) in mIMCD-3 cells, suggesting that 1 µM **TGP-21-RiboTACRiboTAC** might rescue the metabolic alterations caused by miR-21 overexpression in ADPKD, *vide infra*.

In addition to de-repressing PPARα, **TGP-21-RiboTAC** also boosted levels of PDCD4 protein levels by 110 ± 36% (p < 0.05) in WT-9-7 cells and by 150 ± 47% (p < 0.05) in mIMCD-3 cells (**Fig. 5B-C**), as determined by Western blotting. No effect was observed at the *PDCD4* transcript level in either cell lines (**Supplementary Fig. 3A-B**), indicating miR-21 translationally represses *PDCD4* mRNA rather than induces its cleavage.

**Figure 5:**
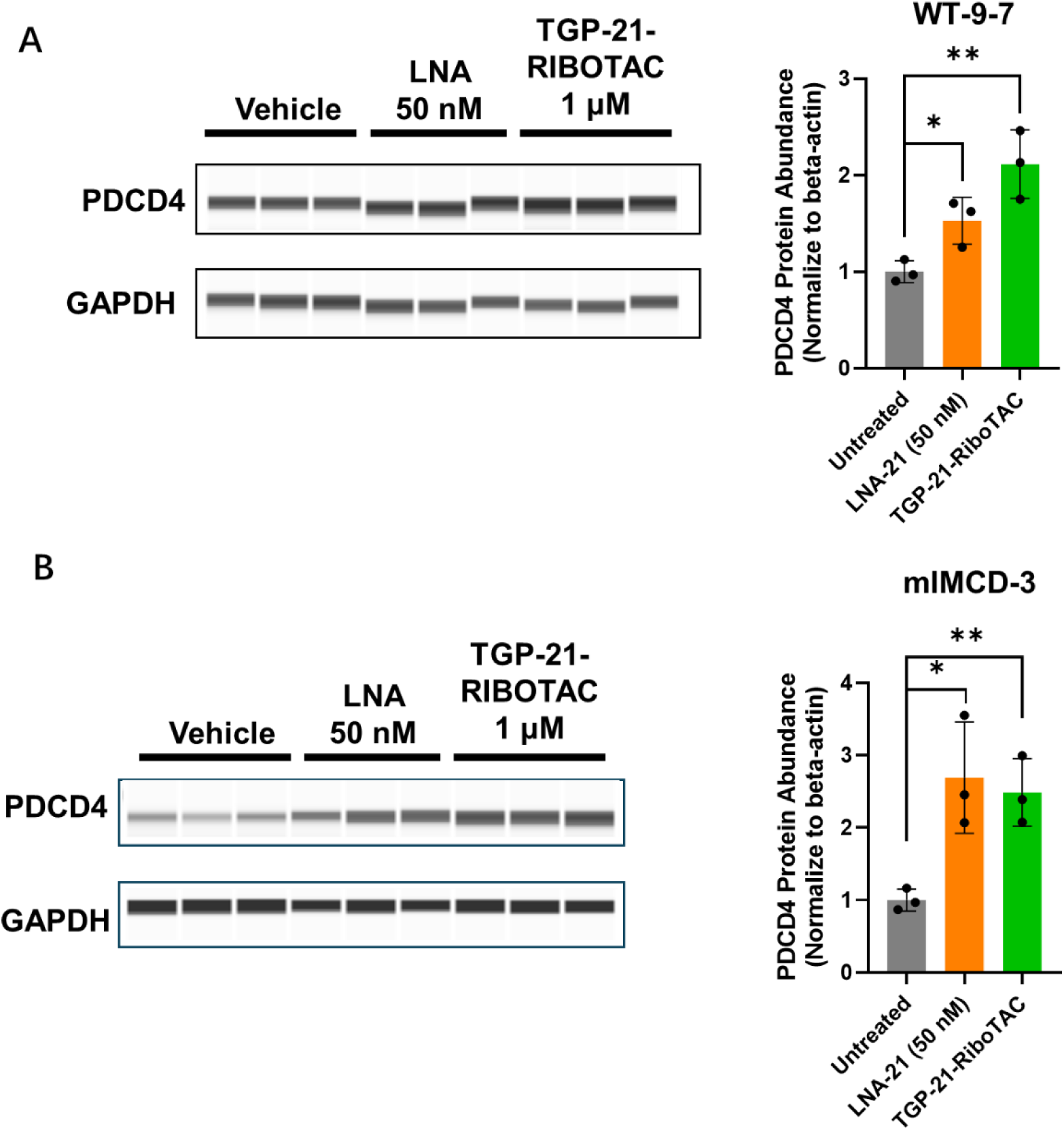
TGP-21-RiboTAC de-represses PDCD4. **A,** Representative Western blot and quantifiction thereof of PDCD4 protein expression in WT-9-7 cells upon **TGP-21-RiboTAC** treatment (n = 3 biological replicates). **B,** Representative Western blot and quantifiction thereof of PDCD4 protein expression in mIMCD-3 cells upon **TGP-21-RiboTAC** treatment (n = 3 biological replicates). *, p < 0.05; **, p < 0.01; ***, p < 0.001; and ****, p < 0.0001, as determined by t tests. Data are reported as mean ± SD.

### TGP-21-RiboTAC reverses ADPKD cellular phenotypes

De-repression of miR-21’s downstream targets PPARα and PDCD4, suggested that **TGP-21-RiboTAC** might rescue ADPKD-associated cellular phenotypes, including apoptosis, proliferation, and cyst formation^2, 3, 11, 14, 16^. The effect of **TGP-21-RiboTAC** on the proliferation of WT 9-7 cells was measured and compared to HEK293T cells, which expresses very low levels of miR-21^28^. These cells serve as a negative control cell line where **TGP-21-RiboTAC** should have no effect. Indeed, a dose dependent reduction in proliferation was observed after treatment of WT 9-7 cells with **TGP-21-RiboTAC** for 2 days and 4 days, where a maximum reduction of 31 % (p < 0.0001) was observed at the 1 µM dose on Day 4 **(Fig. 6A)**. Importantly, no effect was observed in HEK-293T cells at any dose at either time point **(Fig. 6A)**. A locked nucleic acid (**LNA**) antagomir targeting miR-21 was used as a control in both cell lines, reducing the proliferative phenotype in WT 9-7 but not HEK293T cells (**Fig. 6A**). Complementarily, to determine if the observed de-repression was sufficient to induce apoptosis, Caspase 3/7 activity was measured as a proxy ^29, 30^. Apoptosis was increased by 33 ± 11% (p < 0.0001) and 41 ± 11% (p < 0.0001) in WT-9-7 cells, meanwhile by 80 ± 5% (p < 0.0001) and 133 ± 5% (p < 0.0001) in mIMCD-3 cells, upon treatment with 0.5 µM and 1 µM of **TGP-21-RiboTAC**, respectively (**Fig. 6B**).

**Figure 6:**
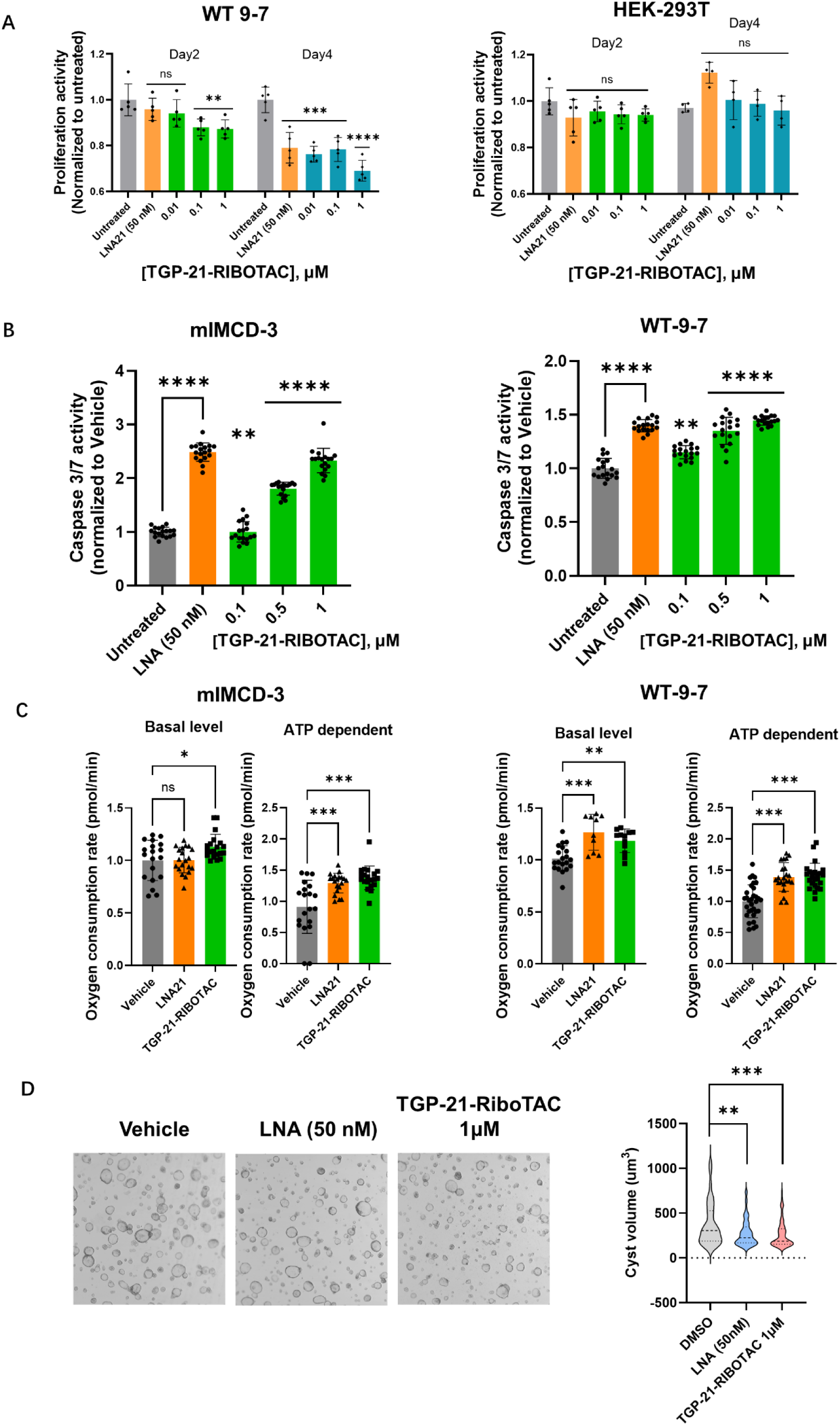
TGP-21-RiboTAC inhibits cell proliferation, cyst growth, and rescues metabolic defects in polycystic kidney cell lines. **A,** Inhibitory effect of **TGP-21-RiboTAC** (0.01 µM, 0.1 µM, 1 µM) and a miR-21-targeting LNA antagomir (50 nM) on the proliferation of WT-9-7 and HEK-293T cells. HEK-293T cells do not express miR-21 and thus should not be affected by RiboTAC-treatment. **B**, Effect of **TGP-21-RiboTAC** on Caspace 3/7 activity, a proxy for apoptosis, in mIMCD-3 cells and WT-9-7 cells. (n = 18 biological replicates). **C,** Quantification of oxygen consumption rate at basal level and ATP dependent mode in mIMCD-3 and WT-9-7 cells upon **TGP-21-RiboTAC** treatment. **D,** Representive images and quantification of cyst growth in mIMCD-3 cells upon **TGP-21-RiboTAC** treatment. (n = 70 biological replicates). *, p < 0.05; **, p < 0.01; ***, p < 0.001; and ****, P < 0.0001, as determined by t tests. Data are reported as mean ± SD.

As previously noted, changes in ATP-dependent oxygen consumption rates^25^ and cyst formation ^31^ are phenotypic defects observed in polycystic kidney disease. Using a Seahorse assay system^32^, the oxygen consumption rate at both basal levels and ATP-dependent levels in mIMCD-3 and WT-9-7 cell line were measured. In the mIMCD-3 cell line, **TGP-21-RiboTAC** increased oxygen consumption rate by 13 ±3% (p < 0.05) at the basal level and 38 ± 4% (p < 0.005) at the ATP-dependent level. Similarly, in WT-9-7 cells, oxygen consumption rate was increased by 19 ± 3% (p < 0.01) at the basal level and 40 ± 4% (p < 0.005) at the ATP-dependent levels. (**Fig. 6C**). As the decreased oxygen consumption rate in mitochondria has positive correlation with increased cyst formation, we next evaluated whether this reduction was sufficient to repress cyst formation in ADPKD cells. To promote cyst formation in mIMCD-3 cells, a 3D cell culture environment was created using Matrigel and induction of cyst formation was achieved by Forskolin treatment. After 4 days of treatment, high content imaging and subsequent quantification revealed that treatment with **TGP-21-RiboTAC** reduced the average cyst volume by 30 ± 4% (p < 0.001) (**Fig. 6D**).

### TGP-21-RiboTAC deactivates the miR-21-SMAD7 circuit in a lung fibroblast cell line

As overexpression of miR-21 is also implicated in lung fibrosis, the activity of **TGP-21-RiboTAC (**1 µM ) was also assessed in retarding the transition of MRC-5 lung fibroblasts to myofibroblasts (**Fig. 7A**) in response to TGF-β ^33, 34^ The abundance of pre-miR-21 and miR-21-5p were measured at 0 h, 12 h, 24 h, and 48 h after addition of TGF-β. In both **TGP-21-RiboTAC**- and vehicle-treated samples, pre-miR-21 and miR-21-5p expression increased as a function of time (**Fig. 7B**). However, levels in **TGP-21-RiboTAC**-treated samples were significantly lower compared to vehicle (**Fig. 7B**). The largest effect was observed 24 h post-TGF-β treatment, where pre-miR-21 and miR-21-5p levels were reduced by 31 ± 4% (p < 0.01) and 39 ± 5% (p < 0.01) in RiboTAC-treated samples, respectively (**Fig. 7B**), and were similar to the levels in vehicle group before TGF-β treatment. At 0 h, 12 h, and 48 h timepoints, miR-21 levels in **TGP-21-RiboTAC** treated samples were 33 ± 6% (p < 0.01), 3 ±17% (p < 0.05) and 29 ± 6% (p < 0.01) lower than the corresponding vehicle-treated samples, and pre-miR-21 levels in **TGP-21-RiboTAC** treated samples were 30 ± 8% (p < 0.05), 24 ± 9% (p < 0.05) and 21 ± 6% (p < 0.05) lower than the corresponding vehicle-treated samples, respectively.

**Figure 7:**
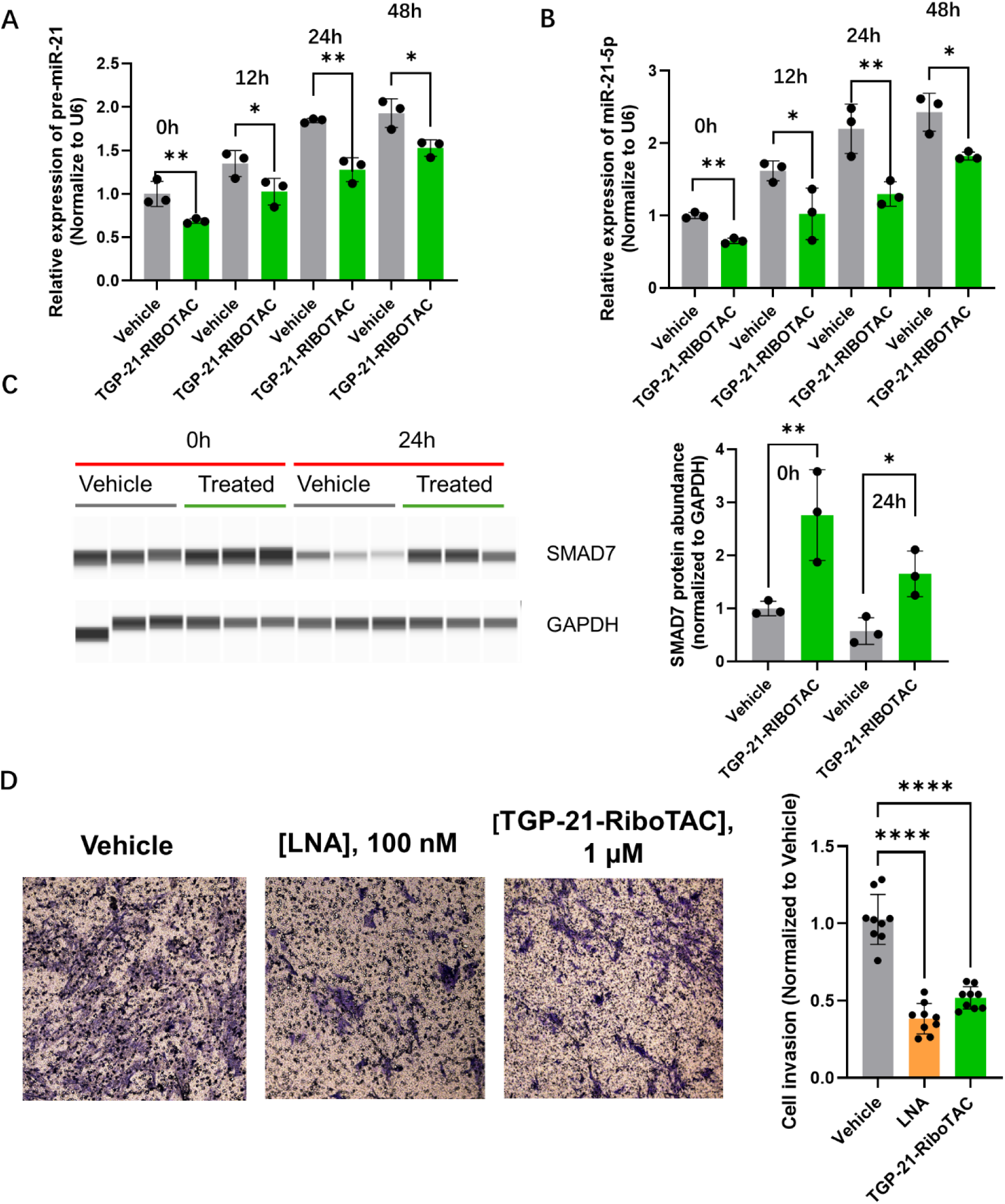
Evaluation of TGP-21-RiboTAC in lung fibroblast cell line MRC-5. **A, Effect of TGP-21-RiboTAC** on pre-miR-21 abundance in MRC-5 cells as a function of time post TGF-β stimulation. **B,** Effect of **TGP-21-RiboTAC** on miR-21-5p abundance in MRC-5 cells as a function of time post TGF-β stimulation. **C.** Representative Western blot and quantifiction of SMAD7 protein in MRC-5 cell line upon **TGP-21-RiboTAC**-treatment. (1 μM; 24 h treatment period; n = 3 replicates per treatment group). **D,** Left: representative images of a Boyden chamber assay to assess migration of MRC-5 cells upon treatment with **TGP-21-RiboTAC** (1 μM) and a miR-21-targeting **LNA** (100 nM) (Cotreatment with TGF-β). Right: quantification of the migration assay (n = 9 biolgoical replicates; 3 images analyzed per biological replicate). *, p < 0.05; **, p < 0.01; ***, p < 0.001; and ****, p < 0.0001, as determined by t tests. Data are reported as mean ± SD.

SMAD7, a direct target of miR-21^9^, is a regulatory protein that acts as a negative regulator of the TGF-β signaling pathway, and plays a crucial role in cell growth and differentiation (**Fig. 7C**)^35, 36^. Therefore, the abundance of SMAD7 in MRC-5 cells upon **TGP-21-RiboTAC** with or without TGF-β stimulation was measured by Simple Western. In the absence of TGF-β stimulation, **TGP-21-RiboTAC** treatment (1 µM) increased the SMAD7 protein expression by 1.8 ± 0.9-fold (p < 0.001) compared to vehicle. As expected, the addition of TGF-β reduced SMAD7 protein abundance in both vehicle- and **TGP-21-RiboTAC**-treated samples, but to a much lesser extent in the latter. SMAD7 levels in MRC-5 myofibroblasts treated with **TGP-21-RiboTAC** increased 1.9 0.8-fold (p < 0.05) compared with vehicle-treated samples (**Fig. 7C**).

As TGF-β signaling induces an invasive phenotype^37^, we studied whether treatment with 1 µM of **TGP-21-RiboTAC** could reduce migration of MRC-5 myofibroblasts (differentiation stimulated by TGF-β). After a 48 h treatment period. **TGP-21-RiboTAC** (1 µM) reduced cellular migration by 51 ± 6% (p < 0.001), similar to treatment with 100 nM of a miR-21-targeting LNA antagomir (64 ± 6%; p < 0.001; **Fig. 7D**). Together, these results demonstrate **TGP-21-RiboTAC** decreases miR-21-5p levels, de-represses SMAD7, and inhibits cellular migration in lung fibrosis.

## DISCUSSION

Aberrant expression of miRNAs contributes to the pathogenesis of a wide spectrum of diseases, yet therapeutic strategies to directly modulate these non-coding RNAs remain limited. In this work, we demonstrate that **TGP-21-RiboTAC**, a small molecule designed to recruit RNase L to its RNA target, selectively degrades the precursor of miR-21 and restores normal gene expressions in models of fibrotic disease. By chemically assembling the endogenous RNase L machinery onto pre-miR-21, this compound reduces mature miR-21 levels, leading to de-repression of downstream targets such as PDCD4, PPARα, and SMAD7. These molecular effects translate into phenotypic rescue across kidney and lung fibrosis cell models—manifested as decreased cyst growth, restoration of mitochondrial metabolism, and inhibition of fibroblast invasion. Together, these findings establish a broadly applicable small-molecule strategy for targeted degradation of disease-causing miRNAs.

The results presented here extend the concept of RiboTAC-mediated RNA degradation beyond oncology into fibrotic disorders such as ADPKD and pulmonary fibrosis, both of which are driven by persistent activation of TGF-β and metabolic stress pathways. Pharmacologic degradation of pre-miR-21 by **TGP-21-RiboTAC** interrupts this fibrotic signaling network at its origin—the miRNA biogenesis stage—rather than at downstream effectors. This mode of action differs fundamentally from oligonucleotide-based antagomirs or inhibitors, as it leverages a catalytic, enzyme-recruiting mechanism that amplifies RNA turnover with high selectivity and does not rely on stoichiometric binding. The restoration of PPARα-dependent metabolism and PDCD4-mediated apoptosis in kidney cells, alongside the normalization of SMAD7-TGF-β signaling in lung fibroblasts, demonstrates the versatility of this approach across distinct tissue contexts.

Collectively, this study provides proof-of-concept that small molecules can be rationally designed to degrade specific non-coding RNAs through endogenous nucleases, offering a new therapeutic paradigm for diseases driven by dysregulated RNA expression. The ability to tune RNA degradation via chemical design opens opportunities to target pathogenic miRNAs and other structured RNAs previously considered “undruggable.” Given the broad role of miR-21 in fibrosis, cancer, and inflammation, **TGP-21-RiboTAC** represents both a chemical probe for dissecting miRNA biology and a lead scaffold for developing small molecule therapeutics that modulate RNA function with precision.

## FIGURES

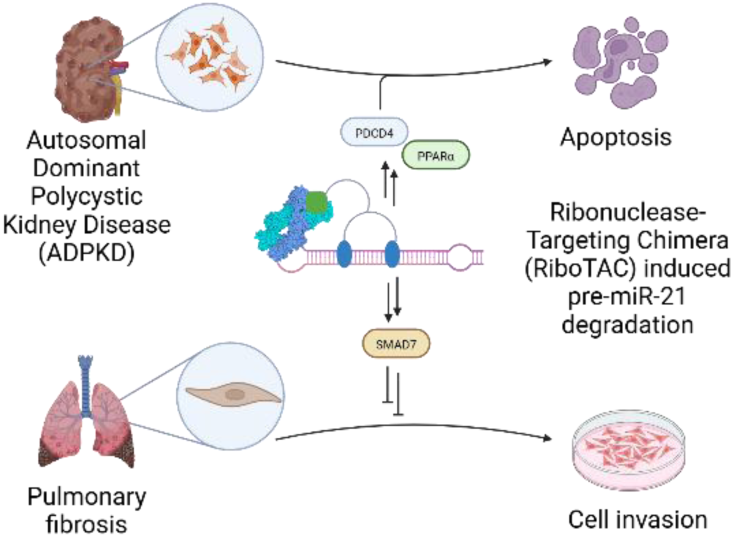

