## Supplemental Information for "Small Molecule Degradation of the microRNA-21 Precursor Rescues Pathogenic Pathways in Cellular Models of Fibrosis"

### EXTENDED METHODS

### GENERAL METHODS

The sequences of all RNA oligonucleotides are provided in Supplemental Table S1. DNA oligonucleotides were purchased from Integrated DNA Technologies (IDT), Inc. with standard desalting and were used without further purification.

**TGP-21** and **TGP-21-RiboTAC** were synthesized and characterized using previously reported procedures[1, 2].

### CELLULAR METHODS

**Cell Culture.** HEK293T cells (CRL-3216) were acquired from ATCC and were maintained in Dulbecco's Modified Eagle Medium (DMEM; Corning, catalog 15-017-CV) supplemented with 10% (v/v) fetal bovine serum (FBS; Sigma Aldrich, catalog #A36707-01), 1% (v/v) Penicillin-Streptomycin Solution (Corning, catalog #30-002-CI) and 1% (v/v) glutaGRO supplement (Corning, catalog #25-015-CI) at 37 °C and 5% $CO_2$.

WT-9-7 cells (CRL-2830) were acquired from ATCC and were maintained in Dulbecco's Modified Eagle Medium (DMEM; Corning, catalog 15-017-CV) supplemented with 10% (v/v) FBS (Sigma Aldrich, catalog #A36707-01), 1% (v/v) Penicillin-Streptomycin Solution (Corning, catalog #30-002-CI) and 1% (v/v) glutaGRO supplement (Corning, catalog #25-015-CI) at 37 °C and 5% $CO_2$. The plates and dishes for WT-9-7 cell culture were pre-incubated with Bovine Collagen Type I Solution (supplied stock concentration is 3.0 mg/mL; PureCol® Advanced Biomatrix Catalog No. 5005-B) for 1 h at 37 °C.

mIMCD-3 cells (CRL-2123) were acquired from ATCC and were maintained in DMEM:F12 Medium (gibco, catalog #11330-32), supplemented with 10% (v/v) FBS (Sigma Aldrich, catalog #A36707-01), 1% (v/v) Penicillin-Streptomycin Solution (Corning, catalog #30-002-CI) and 1% (v/v) glutaGRO supplement (Corning, catalog #25-015-CI) at 37 °C and 5% $CO_2$.

MRC-5 cells (ATCC, catalog #CCL-171) were maintained in Minimum Essential Medium Eagle (Sigma-Aldrich, catalog No. M4655) supplemented with 10% (v/v) FBS (Sigma Aldrich, catalog #A36707-01), 1% (v/v) Penicillin-Streptomycin Solution (Corning, catalog No.30-002-CI) and 1% (v/v) glutaGRO supplement (Corning, catalog #25-015-CI) at 37 °C and 5% $CO_2$.

All cell lines were seeded into 6-well or 12-well plates and allowed to grow to ~40% confluency for compound treatment. Growth medium was removed and replaced with fresh medium containing the compound of interest (DMSO < 0.5% (v/v)). After 48 h, the medium was removed, and the cells were washed with 1× DPBS and then harvested for downstream analysis as described for each experiment type.

**Measuring Abundance of miRNA and mRNA by RT-qPCR**. Cells were plated in 12-well plates and grown to ~50% confluency. Following compound treatment, total RNA was extracted using a Quick-RNA Miniprep Kit (Zymo Research, catalog #D4069) following the manufacturer's protocol, including the on-column DNase I digestion. RNA concentration was determined using a Nanodrop UV spectrophotometer (ThermoFisher), only samples with $OD_{260}/OD_{280}$ >1.8, were further processed.

For measuring mature miRNA abundance by RT-qPCR, approximately 200 ng of total RNA was used for reverse transcription (RT) using a Taqman MicroRNA Reverse Transcription

Kit (Applied Biosystems, catalog # 4366596) per manufacturer's protocol. Next, 20 ng of cDNA from the RT reaction was supplemented with primers and Taqman Universal PCR Master Mix (Applied Biosystems, catalog # 4324018). To measure pre-miR-21 abundance, 200ng of total RNA was used for reverse transcription (RT) using a qScript cDNA synthesis kit (Quantabio, catalog # 95047) per manufacturer's protocol. Next, 20 ng of cDNA from the RT reaction was supplemented with forward and reverse primers (Supplemental Table S4) and SYBR Green Universal Master Mix (Applied Biosystems, catalog #4309155). qPCR amplification was completed using an Applied Biosystems QuantStudio5 instrument. Expression levels of miRNAs were normalized to U6.

For mRNA RT-qPCR, 200ng of total RNA was used for reverse transcription (RT) using a qScript cDNA synthesis kit (Quantabio, catalog # 95047) per manufacturer's protocol. Next, 20 ng of cDNA from the RT reaction was supplemented with forward and reverse primers (Supplemental Table S4) and SYBR Green Universal Master Mix (Applied Biosystems, catalog #4309155). qPCR amplification was completed using an Applied Biosystems QuantStudio5 instrument. Expression levels of mRNAs were normalized to GAPDH.

All RT-qPCR data were analyzed using the ΔΔCt method[3]. See Supplementary Table 4 for a list of primers.

**siRNA Knockdown of RNase L.** WT-9-7 cells were seeded into 12-well plates and grown ~40% confluency before transfection and compound treatment. The RNase L-targeting siRNA-2 (SMARTpool ON-TARGET plus) and control siRNA-2 (ON-TARGET plus) were purchased from Dharmacon. The siRNA of interest (100 nM) was transfected with Lipofectamine RNAiMAX reagent (Invitrogen, catalog #13778150) per manufacturer's protocol.

After the transfection, DMSO or **TGP-21-RIBOTAC** (1 μM) was added to cells, and the cells were treated for 48 h. Total RNA was then harvested, and RT-qPCR was performed as described above.

**Simple Western to Measure Protein Abundance.** Cells were cultured in 6-well plates until they reached ~50% confluency. They were then treated as described in "Cell Culture". Total protein was extracted using 200 μL of Mammalian Protein Extraction Reagent (MPER; ThermoScientific) with 1× Protease Inhibitor Cocktail III (RPI P50700-1) per the manufacturer's protocol. Protein concentration was quantified by using a Pierce BCA kit (catalog #23255) in 96-well plate per manufacturer's protocol. To measure endogenous PDCD4, PPARα, or SMAD7 protein abundance, a Simple Western was completed using a Jess Automated Western Blot System (Protein Simple) per the manufacturer's protocol. A 14 μL aliquot of 0.4 mg/mL total protein was loaded into the Simple Western instrument. The β-Actin primary antibody (Cell Signaling Technology, catalog #8H10D10), PDCD4 primary antibody (Cell Signaling Technology, catalog #D29C6), PPARα primary antibody (Cell Signaling Technology, catalog #7076P2), and SMAD7 primary antibody (Santa Cruz, catalog #sc-365846) were each diluted 1:100. Secondary antibody (Anti-Mouse or Anti-Rabbit) was provided by manufacturers without any dilution. Chemiluminescence mode and twenty-five capillary cartridges were used. Data are reported as the ratio of PDCD4, PPARα, or SMAD7 to β-Actin, normalized to vehicle-treated samples.

**Caspase-3/7 Activity.** WT-9-7 cells were cultured in 96-well plates until they reached ~40% confluency. They were then treated as described in "Cell Culture". After a 48 h treatment period, Caspase-3/7 activity was measured using Caspase-Glo® 3/7 Assay System

(Promega, catalog #G8090) per the manufacturer's recommended protocol. Luminescence was measured using a Tecan Infinite M1000 Pro plate reader (integration time =500 ms). Data is normalized to vehicle-treated samples.

**Immunofluorescence (IF) Assay.** WT-9-7 cells were seeded in the glass bottomed, 24-well plate at a density of 50,000 cells/well. Upon reaching ~20% confluency, the cells were either treated with **TGP-21-RiboTAC** (1 μM), LNA-21 (50 nM), or DMSO (0.2% (v/v)). After incubating for 48 h, the growth medium was removed, and cells were washed by 1× DPBS twice. The cells were then fixed with 4% (w/v) paraformaldehyde prepared in 1× DPBS (400 μL per well) at room temperature for 10 min, followed by washing with 500 μL of 1× DPBS twice. The cells were incubated with blocking buffer (1× PBS containing 5% (v/v) FBS, and 0.3% (v/v) TritonTM X-100) for 1 h at room temperature. After removing the blocking buffer, the cells were incubated with PPARα primary antibody solution (PPARα primary antibody (Cell Signaling Technology, catalog #7076P2), diluted 1:500 ratio in Antibody Dilution Buffer (1× PBS supplemented with 1% (w/v) BSA and 0.3% Triton (v/v) X-100)) overnight at 4 °C. The primary antibody solution was removed, and the cells were washed three times with 500 μL of 1× PBS for 5 min each. The cells were then incubated with Alex488-conjugated mouse secondary antibody (Cell Signaling Technology, catalog #4408) diluted 1:1000 in Antibody Dilution Buffer for 1 h at room temperature. After removing the secondary antibody solution, cells were incubated with Hoechst diluted 1:5000 in 1× PBS for 15 min (final concentration of 2 μg/mL) at room temperature. The cells were rinsed three times with 200 μL of 1× PBS for 5 min each and imaged by a confocal microscopy (10× magnification; two views captured per well, Olympus FV3000). Fluorescence intensity of images was quantified with ImageJ.

**Oxygen Consumption Rate (OCR) Assay.** WT-9-7 cells or mIMCD-3 cells were seeded in 12-well plates at ~60% confluency. Cells were treated with **TGP-21-RIBOTAC** (1 μM), LNA-21 (Qiagen, 50 nM), or vehicle at the indicated concentrations for 24 h in growth medium. The cells were then trypsinized and seeded into an XF96 cell culture microplate (Seahorse Bioscience) (mIMCD-3 (5,000 cells/well); WT-9-7 (4,000 cells/well). Cells were treated again as indicated and incubated for an additional 24 h. Mitochondrial OCRs were determined using an XF96 extracellular flow analyzer (Seahorse Bioscience) following the manufacturer's protocol. In brief, mitochondrial respiration rate was measured with a basal-oligomycin-carbonyl cyanide 4-(trifluoromethoxy) phenylhydrazone (FCCP)-antimycin-A/rotenone 'BOFA' experiment[4, 5],. After incubation, cells were washed with 100 μL of 1× DPBS. Cells were then equilibrated for 1 h at 37 °C (ambient $CO_2$) in XF Assay DMEM Medium (Seahorse Bioscience) (pH 7.4), supplemented with 1 mM sodium pyruvate, 1 mM L-glutamine, and 7 mM glucose. The XF96 plate was then transferred to a temperature-controlled (37 °C) Seahorse analyzer. To measure the basal rate, cells were subjected to a 10 min equilibration period and three assay cycles, comprising a 3 min mix, a 2 min wait, and a 3 min measure period, without adding any substrates. To obtain the ATP-dependent OCR, cells were subjected to a 10 min equilibration period and oligomycin (ATP synthase inhibitor, 1 μM final concentration) was then added by automatic pneumatic injection followed by three assay cycles, comprising of 3 min mix, 2 min wait and a 3 min measure period. After running the assay, cells were stained with Hoechst immediately and counted by Cytation 5 plate reader (Agilent Technologies). Measured OCR was normalized with viability of cell in each well. The

ATP-dependent OCR was calculated by subtracting the OCR reading recorded after oligomycin treatment from OCR reading recorded at basal levels.

ATP-linked OCR = (basal OCR - post-Oligomycin OCR)/ Normalized cell viability (measured with Hoechst)

**Cyst Formation Assay.** A 200 μL aliquot of Matrigel matrix (Corning, catalog #356234) (8 to 11 mg/mL) was added into each well of a pre-chilled (4 °C) 24-well plate, and the matrix was spread evenly with a pipet tip. The plate was then incubated at 37 °C for 30 min to allow the matrix to form a gel. mIMCD-3 cells were washed once with PBS and isolated from the plate using trypsin to make a single-cell suspension; cells were pelleted through centrifugation at 125 × g for 5 min at room temperature. The pelleted cells were re-suspended in mIMCD-3 complete medium (DMEM:F12 Medium + 10% (v/v) FBS) with 100 nM forskolin (Sigma Aldrich, catalog #F6886) to adjust the final cell density to $3 \times 10^5$ cells/mL, and 250 μL of the cell suspension was added into each well of the Matrigel-coated 24-well plate. The cells were incubated at 37 °C for 30 min. mIMCD-3 complete medium supplemented with 100 nM forskolin was chilled on ice and added to Matrigel matrix to make a 10-fold dilution of Matrigel matrix final concentration: 0.8 to 1.1 mg/mL). A 250 μL aliquot of the Matrigel matrix-medium mixture was then gently added to the plated culture. Cells were cultured for 4 to 7 days, and the Matrigel matrix-medium mixture (with 100 nM forskolin) was changed every 2 days.

Images were analyzed in R using the EBImage[6] package with a customed script. Raw TIFF images were loaded and converted to grayscale, and a 3 × 3 high-pass filter (Laplacian kernel) was applied to emphasize local intensity transitions. To avoid border effects, the filtered image was cropped to the central 761 × 761-pixel region. The cropped image was then binarized using adaptive local thresholding (window size 5 × 5 pixels, offset = 0.18), and refined by applying a series of morphological operations including dilation, erosion and hole fill. The binary image was then segmented to isolate each individual "cyst" objects. The pixel area of each labeled object was computed, and objects smaller than 50 pixels were classified as debris and removed. Objects identified across all images from a given condition were pooled into one dataset for statistical analysis of cyst size.

**Cell Proliferation Assay.** Cell proliferation assays were completed using CellTiter 96® AQueous One Solution Cell Proliferation Assay Solution (Promega, catalog #G3580) per manufacturer's protocol. WT-9-7 or HEK293T cells were seeded into 96-well white, clear bottom tissue culture plates (Corning, catalog #3610). Upon reaching ~15% confluency, the cells were either treated with **TGP-21-RiboTAC** (1 μM), LNA-21 (50 nM), or DMSO (0.2% (v/v)). After incubation for the indicated times (2 or 4 days), 20 μL of the proliferation assay solution was added to each well, and the samples were incubated at 37 °C for 30 min. The absorbance at 490 nm was then measured using a SpectraMax M5 fluorescence plate reader. The results were normalized to vehicle treated samples.

**Boyden Chamber Invasion Assay.** Hanging cell culture inserts (8.0 μm; for 24-well plates) were precoated with Matrigel (0.3 mg/mL; Corning 356234). MRC-5 cells (75,000 per insert) were seeded into the hanging inserts in serum-free growth medium containing 8 ng/mL TGF-β (100 μL), with or without compound (0.1% (v/v) final DMSO concentration). The inserts were placed into 24-well plates containing 650 μL of complete growth medium containing 8 ng/mL TGF-β. After 48 h incubation at 37 °C, the medium in the insert was removed and washed with 600 μL of 1× DPBS. The cells were then fixed with 600 μL of 4%

(w/v) paraformaldehyde in 1× DPBS for 20 min at room temperature. The inserts were washed twice with 1× DPBS (600 μL each) and stained with 600 μL of 0.1% (w/v) crystal violet in 20% methanol (v/v) for 20 min. After washing twice with Nanopure water (600 μL each) and once with 1× DPBS (600 μL), noninvasive cells on the surface of the Matrigel were removed with cotton swabs. Invading cells were imaged under a Leica DMI3000 B upright fluorescent microscope with four fields of view assessed per sample. Images were analyzed using imageJ by calculating the ratio of area of invasion cells to area of fields. Basically, the threshold was adjusted to isolate the invaded cells by going to Image > Adjust > Threshold. Next, the ratio of area of invasion cells to fields was measured by going to Analyze > Analyze Particles and setting the size range (300 to infinity here)[7]. Results were normalized to vehicle.

### Supplementary Figures

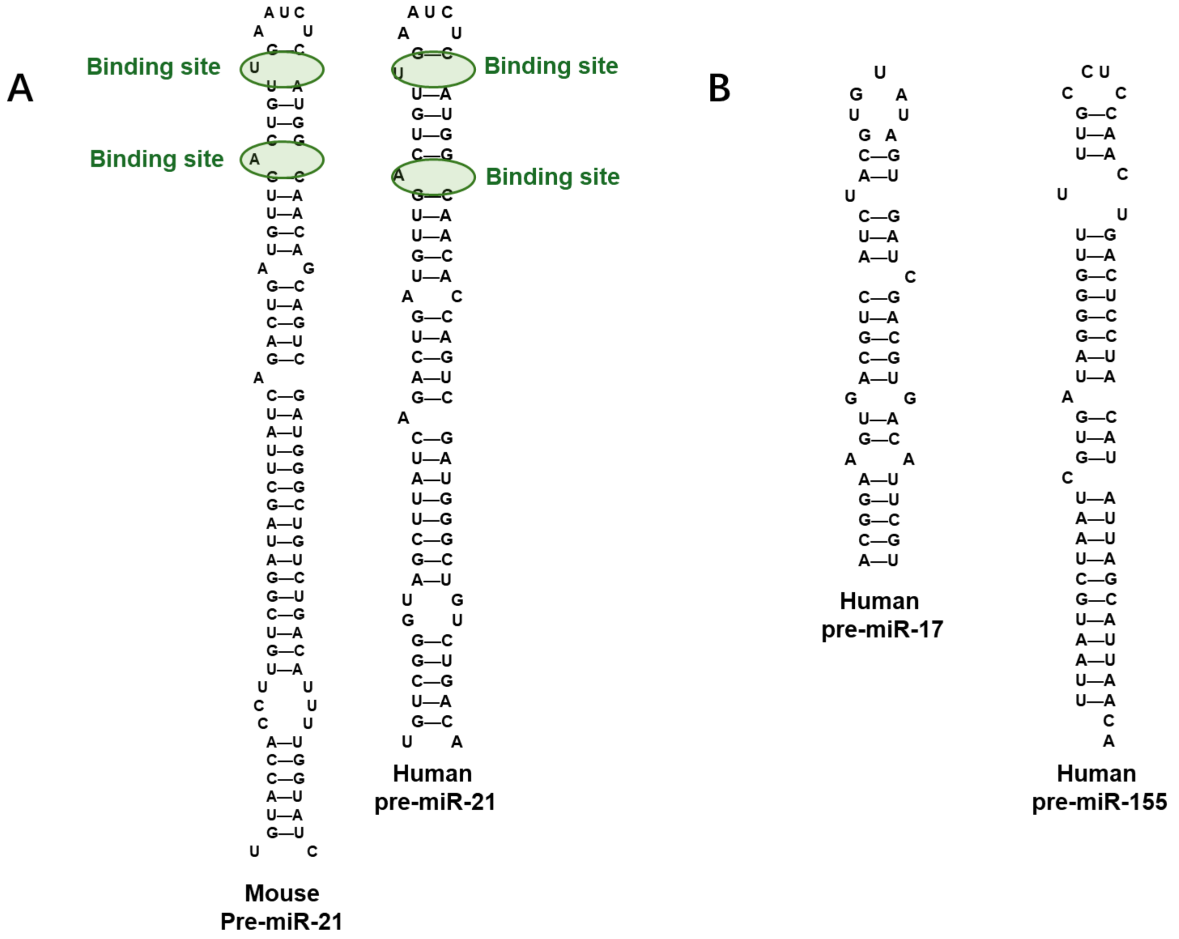


**Fig. S1. Secondary structures of pre-miR21, pre-miR-17 and pre-miR-155. A)** Secondary structures of pre-miR-21 from mouse and human. The binding sites for dimeric **TGP-21** are shown in green circles. **B)** Secondary structures of human pre-miR-17 and pre-miR-155.

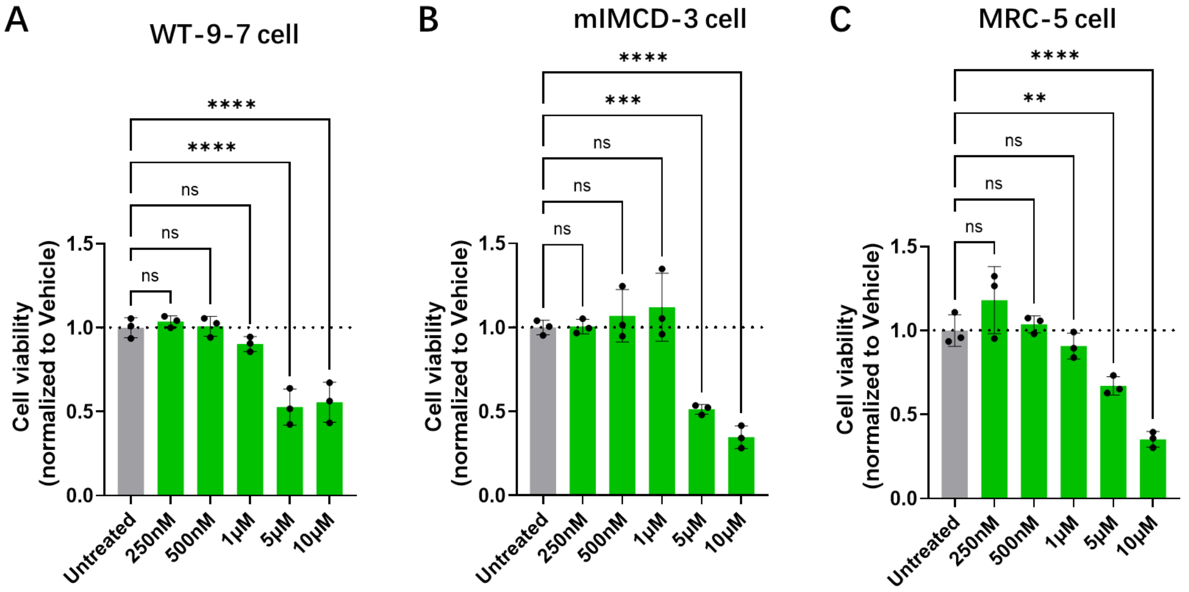


**Fig. S2. Effect of TGP-21-RiboTAC on the viability of in WT-9-7, miMCD3, and MRC-5 cells.** **A**) Effect of **TGP-21-RiboTAC** treatment (250 nM, 500 nM, 1 μM, 5 μM and 10 μM) on the viability of WT-9-7 cells upon for 48 h treatment (n=3 biological replicates). **B**) Effect of **TGP-21-RiboTAC** treatment (250 nM, 500 nM, 1 μM, 5 μM and 10 μM) on the viability of miMCD3 cells upon for 48 h treatment (n=3 biological replicates). **C**) Effect of **TGP-21-RiboTAC** treatment (250 nM, 500 nM, 1 μM, 5 μM and 10 μM) on the viability of MRC-5 cells upon for 48 h treatment (n=3 biological replicates). $p < 0.05$; **, $p < 0.01$; ***, $p < 0.001$; and ****, $p < 0.0001$, as determined by a One-way ANOVA with multiple comparisons. Data are reported as mean ± SD.

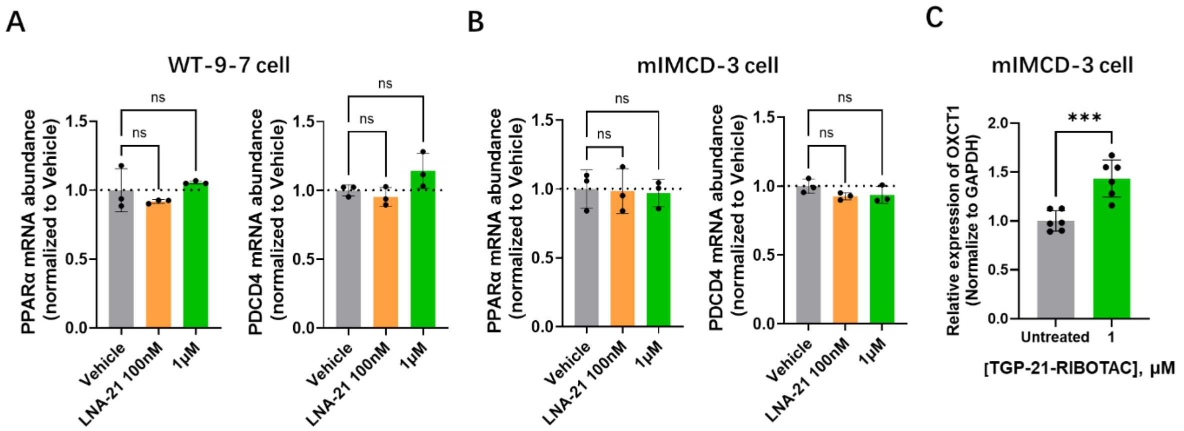


**Fig. S3. Effect of TGP-21-RiboTAC on *PPARα* and *PDCD4* mRNA levels in WT-9-7 and mIMCD-3 cells.** **A**) Abundance of *PPARα* and *PDCD4* mRNA in WT-9-7 cells upon **TGP-21-RiboTAC** treatment (1 μM) for 48 h (n=3 biological replicates). **B**) Abundance of PPARα and *PDCD4* mRNA in mIMCD-3 cells upon **TGP-21-RiboTAC** treatment (1 μM) for 48 h. (n=3 biological replicates). **C)** Effect of **TGP-21-RiboTAC** on PPARα target gene *oxct1* expression in WT-9-7 cells, as determined by RT-qPCR (n=3 biological replicates). $p < 0.05$; **, $p < 0.01$; ***, $p < 0.001$; and ****, $p < 0.0001$, as determined by a One-way ANOVA with multiple comparisons. Data are reported as mean ± SD.

**SUPPLEMENTARY TABLES**

**Table S1. Sequences of oligonucleotides used in this study.** "Fwd" denotes forward primer and "Rev" denotes reverse primer; "h" denotes human and "m" denotes mouse.

| Oligonucleotide | Sequence 5' → 3' or Assay ID | Experiment | Supplier |
|---|---|---|---|
| Pre-miR-21 Fwd | CTGATGTTGACTGTTGAATC | qPCR | IDT |
| Pre-miR-21 Rev | GCCCATCGACTGGTGTTGCC | qPCR | IDT |
| hGAPDH Fwd | TGCACCACCAACTGCTTAG | qPCR | IDT |
| hGAPDH Rev | GATGCAGGGATGATGTTC | qPCR | IDT |
| mGAPDH Fwd | AGGTCGGTGTGAACGGATTTG | qPCR | IDT |
| mGAPDH Rev | TGTAGACCATGTAGTTGAGGTCA | qPCR | IDT |
| Mature miR-21 | 000397 | qPCR | Applied bio |
| U6 | 001973 | qPCR | Applied bio |
| Mature miR-155 | 467534_mat | qPCR | Applied bio |
| Mature miR-17 | 000393 | qPCR | Applied bio |
| hPPARα Fwd | GGAGTTTATGAGGCCATATTCG | qPCR | IDT |
| hPPARα Rev | CAAAATCAAACTTGGGTTCCAT | qPCR | IDT |
| mPPARα Fwd | ACCACTACGGAGTTCACGCATG | qPCR | IDT |
| mPPARα Rev | GAATCTTGCAGCTCCGATCACAC | qPCR | IDT |
| hPDCD4 Fwd | ACTGTGCCAACCAGTCCAAAGG | qPCR | IDT |
| hPDCD4 Rev | CCTCCACATCATACACCTGTCC | qPCR | IDT |
| mPDCD4 Fwd | GCGGTTAGAAGTGGAGTTGCTG | qPCR | IDT |

| | | | |
|---|---|---|---|
| mPDCD4 Rev | CCACATCATACACCTGTCCAGG | qPCR | IDT |
| hSLC27A Fwd | GTGGAGAAAGATGAACCTGTCG | qPCR | IDT |
| hSLC27A Rev | CTGAGCCTTTGCTCCAGCATAG | qPCR | IDT |
| hCPT2 Fwd | GCAGATGATGGTTGAGTGCTCC | qPCR | IDT |
| hCPT2 Rev | AGATGCCGCAGAGCAAACAAGT | qPCR | IDT |
| hCPAM Fwd | TTGTGGCTTGCCTGCTCCTCTA | qPCR | IDT |
| hGPAM Rev | AATCACGAGCCAGGACTTCCTC | qPCR | IDT |
| hOXCT1 Fwd | GCTTTGGTGAAAGCCTGGAAGG | qPCR | IDT |
| hOXCT1 Rev | CTCTACCACTGTGGTTTCTGCAG | qPCR | IDT |
| mOXCT1 Fwd | GAGCGACAGTTCCTTTCTGGTG | qPCR | IDT |
| mOXCT1 Rev | TCCCATACCCTGTGCTGGTGTA | qPCR | IDT |
